# A public vaccine-induced human antibody protects against SARS-CoV-2 and emerging variants

**DOI:** 10.1101/2021.03.24.436864

**Authors:** Aaron J. Schmitz, Jackson S. Turner, Zhuoming Liu, Ishmael D. Aziati, Rita E. Chen, Astha Joshi, Traci L. Bricker, Tamarand L. Darling, Daniel C. Adelsberg, Wafaa B. Alsoussi, James Brett Case, Tingting Lei, Mahima Thapa, Fatima Amanat, Jane A. O’Halloran, Pei-Yong Shi, Rachel M. Presti, Florian Krammer, Goran Bajic, Sean P.J. Whelan, Michael S. Diamond, Adrianus C. M. Boon, Ali H. Ellebedy

## Abstract

The emergence of antigenically distinct severe acute respiratory syndrome coronavirus 2 (SARS-CoV-2) variants with increased transmissibility is a public health threat. Some of these variants show substantial resistance to neutralization by SARS-CoV-2 infection- or vaccination-induced antibodies, which principally target the receptor binding domain (RBD) on the virus spike glycoprotein. Here, we describe 2C08, a SARS-CoV-2 mRNA vaccine-induced germinal center B cell-derived human monoclonal antibody that binds to the receptor binding motif within the RBD. 2C08 broadly neutralizes SARS-CoV-2 variants with remarkable potency and reduces lung inflammation, viral load, and morbidity in hamsters challenged with either an ancestral SARS-CoV-2 strain or a recent variant of concern. Clonal analysis identified 2C08-like public clonotypes among B cell clones responding to SARS-CoV-2 infection or vaccination in at least 20 out of 78 individuals. Thus, 2C08-like antibodies can be readily induced by SARS-CoV-2 vaccines and mitigate resistance by circulating variants of concern.

**One Sentence Summary:** Protection against SARS-CoV-2 variants by a potently neutralizing vaccine-induced human monoclonal antibody.

## Main Text

SARS-CoV-2 is a highly pathogenic coronavirus that first emerged in Wuhan, Hubei province of China in late 2019 (*1*, *2*). The virus quickly spread to multiple continents, leading to the coronavirus disease 2019 (COVID-19) pandemic. To date, SARS-CoV-2 has caused more than 120 million confirmed infections, leading to approximately three million deaths (*3*). The damaging impact of the morbidity and mortality caused by the COVID-19 pandemic has triggered a global effort towards developing SARS-CoV-2 countermeasures. These campaigns led to the rapid development and deployment of antibody-based therapeutics (immune plasma therapy, monoclonal antibodies (mAbs)) and vaccines (lipid nanoparticle-encapsulated mRNA, virus-inactivated, and viral-vectored platforms) (*4*–*8*). The high efficacy of mRNA-based vaccines in particular has raised hope for ending the pandemic (*9*–*11*). However, the emergence of multiple SARS-CoV-2 variants that are antigenically distinct from the early circulating strains used to develop the first generation of vaccines has raised concerns for compromised vaccine-induced protective immunity (*12*–*14*). Indeed, multiple studies have demonstrated that these variants show reduced neutralization *in vitro* by antibodies elicited in humans in response to SARS-CoV-2 infection or vaccination (*15*–*18*). This observation highlights the need for better understanding of the breadth of SARS-CoV-2 vaccine-induced antibody responses and possible adjustments of prophylactic and therapeutic reagents to combat emerging variants.

SARS-CoV-2 entry into host cells is mediated primarily by the binding of the viral spike (S) protein through its receptor-binding domain (RBD) to the cellular receptor, human angiotensin-converting enzyme 2 (ACE2) (*19*). Thus, the S protein is a critical target for antibody-based therapeutics to prevent SARS-CoV-2 virus infection and limit its spread. Indeed, the RBD is recognized by many potently neutralizing monoclonal antibodies (*20*–*27*). Pfizer-BioNTech SARS-CoV-2 mRNA vaccine (BNT162b2) encodes the full-length prefusion stabilized SARS-CoV-2 S protein and induces robust serum binding and neutralizing antibody responses in humans (*9*, *28*). We recently described the S-specific plasmablast and germinal center (GC) B cell responses induced by BNT162b2 vaccination in healthy adults. GC B cells were analyzed in aspirates from the draining axillary lymph nodes of 12 participants after vaccination. We verified the specificity of the GC response by generating a panel of recombinant human mAbs from single cell-sorted S^+^ GC B cells isolated from three participants (*29*). The majority of these vaccine-induced antibodies are directed against the RBD. Here, we assess the capacity of these anti-RBD mAbs to recognize and neutralize recently emerged SARS-CoV-2 variants.

From a pool of S^+^ GC B cell-derived mAbs, we selected 13 human anti-RBD mAbs that bound avidly to the predominantly circulating WA1/2020 D614G SARS-CoV-2 strain referred to hereafter as the D614G strain (*29*, *30*). We assessed mAbs binding to recombinant RBDs derived from the D614G strain and three SARS-CoV-2 variants, B.1.1.7, B.1.351 and B.1.1.248 by enzyme-linked immunosorbent assay (ELISA). Only one mAb, 1H09, showed decreased binding to the RBD derived from the B.1.1.7 variant (**Fig. 1A**). Four additional mAbs completely lost or showed substantially reduced binding to the B.1.351 and B.1.1.248 variant RBDs (**Fig. 1A**). The remaining eight mAbs showed equivalent binding to RBDs from all tested strains (**Fig. 1A**). We next examined the *in vitro* neutralization capacity of the 13 mAbs against the D614G SARS-CoV-2 strain using a high-throughput focus reduction neutralization test (FRNT) with authentic virus (*31*). Only five mAbs (2C08, 1H09, 1B12, 2B06, and 3A11) showed high neutralization potency against D614G with 80% neutralization values of less than 100 ng/mL. We then assessed the ability of these five mAbs to neutralize the B.1.1.7, B.1.351 and B.1.1.248 variants. Consistent with the RBD binding data, 1H09 failed to neutralize any of the emerging variants, whereas 1B12, 2B06 and 3A11 neutralized B.1.1.7 but not the B.1.351 and B.1.1.248 variants (**Fig. 1B**). One antibody, 2C08, neutralized the four SARS-CoV-2 strains we tested with remarkable potency (half-maximal inhibitory concentration of 5 ng/mL) (**Fig. 1B**), indicating that it recognizes RBD residues that are not altered in these variants.

**Figure 1.**
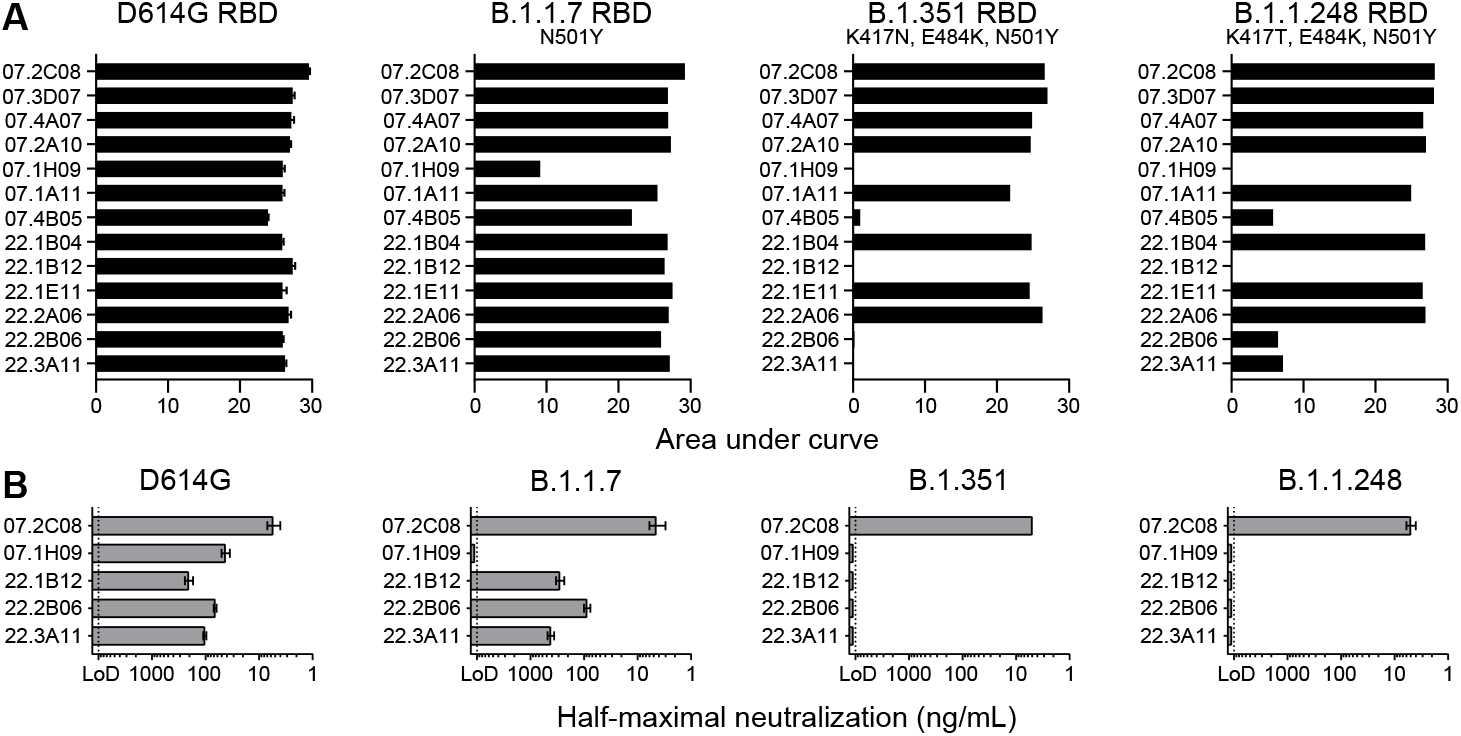
mAb 2C08 potently neutralizes diverse SARS-CoV-2 strains. (**A and B**) ELISA binding to recombinant RBD from (A) and neutralizing activity in Vero-TMPRSS2 cells against (B) indicated SARS-CoV-2 strains by the indicated mAbs. ELISA binding to D614G RBD previously reported in (*29*). Baseline for area under the curve was set to the mean + three times the standard deviation of background binding to bovine serum albumin. Dotted lines indicate limit of detection. Bars indicate mean ± SEM. Results are from one experiment performed in duplicate (panel A, D614G) or in singlet (panel A, B.1.1.7, B.1.351, and B.1.1.248), or two experiments performed in duplicate (panel B).

To assess the protective capacity of 2C08 *in vivo*, we utilized a hamster model of SARS-CoV-2 infection (*32*). We evaluated the prophylactic efficacy of 2C08 against the D614G strain and against a fully infectious recombinant SARS-CoV-2 with B.1.351 spike gene (Wash SA-B.1.351; D80A, 242-244 deletion, R246I, K417N, E484K, N501Y, D614G and A701V) (*16*) in 4–6-week-old male Syrian hamsters. Animals treated with 2C08 and challenged with either virus did not lose weight during the experiment and started to gain weight (relative to starting weight) on 3 dpi. In contrast, animals treated with the isotype control mAb started losing weight on 2 dpi (**Fig 2A**). The average weights between the isotype− and 2C08-treated animals differed by 5.9 percent on 3 dpi (*P* = 0.008) and 7.7 percent 4 dpi (*P* = 0.008) for the D614G challenge and by 6.8 percent on 3 dpi (*P* = 0.095) and 9.1 percent 4 dpi (*P* = 0.056) for the B.1.351 challenge. Consistent with the weight loss data, 2C08 treatment reduced viral RNA levels by more than 10,000-fold in the lungs of the D614G challenged hamsters and by approximately 1000-fold in those challenged with B.1.351 SARS-CoV-2 (*P* = 0.008 for both) (**Fig. 2B, Fig. S1A**) on 4 dpi compared to the isotype control mAb groups. Prophylactic treatment also significantly reduced infectious virus titers for both strains detected in the lungs on 4 dpi (*P* = 0.008 for both) (**Fig 2C**). In addition to viral load, concentrations of proinflammatory cytokines were significantly reduced in animals that received 2C08 prophylaxis (**Fig. 2D**). In comparison to control mAb treated animals, a significant decrease in host gene-expression was observed for *Ccl3, CcL5, Ifit3, Ifit6, Ip10, Irf7 and Rig-I* in lungs of 2C08-treated animals. Overall, prophylaxis with 2C08 showed substantial capacity to decrease viral infection in lower respiratory tissues upon challenge with SARS-CoV-2 strains with spike genes corresponding to ancestral and a key emerging variant.

**Figure 2.**
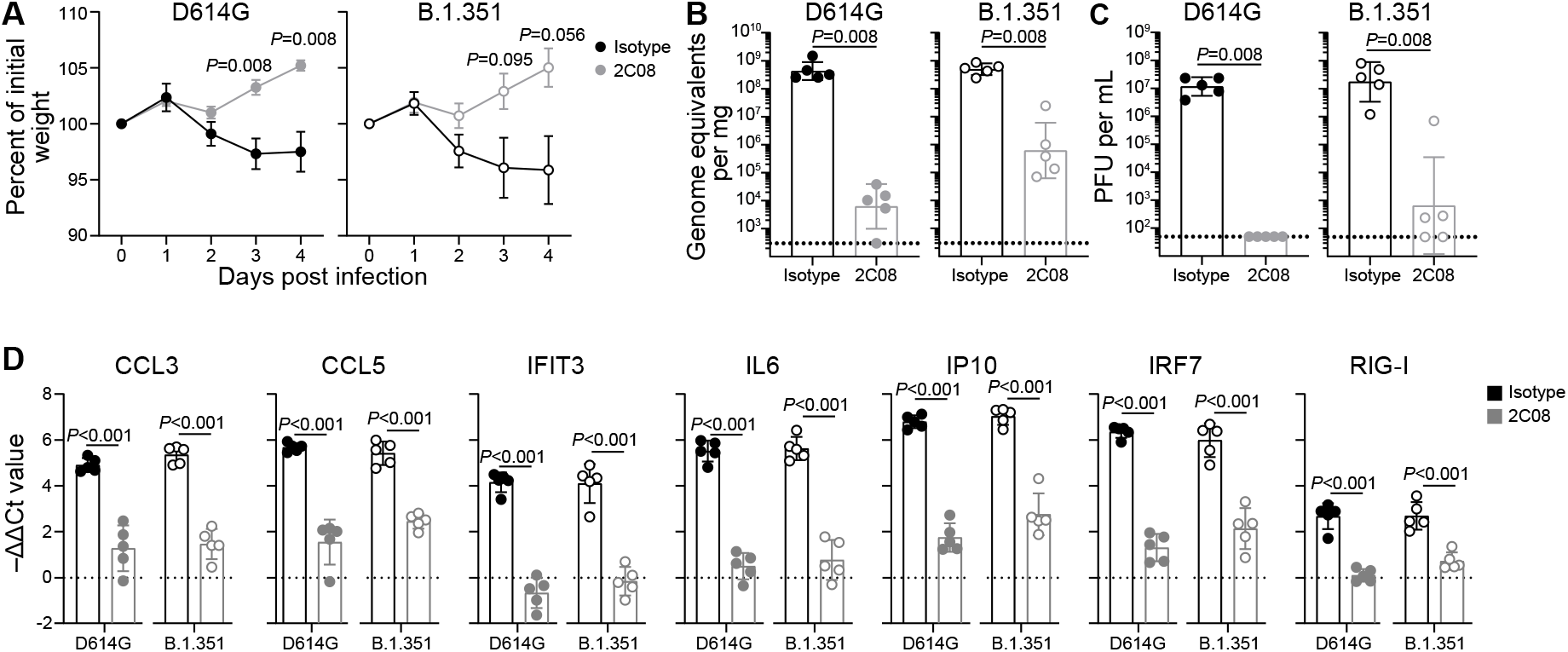
mAb 2C08 protects hamsters from SARS-CoV-2 challenge. **(A–D**) Percent weight change (A), lung viral RNA titer (B), lung infectious virus titer (C), and lung cytokine gene expression (D) of hamsters that received isotype (black) or 2C08 (grey) one day prior to intranasal challenge with 5×10^5^ PFU D614G (*left*) or B.1.351 (*right*) SARS-CoV-2. In (A), symbols indicate mean ± SEM. In (B and C), bars indicate geometric mean ± geometric SD, and each symbol represents one hamster. In (D), bars indicate mean ± SD, and each symbol represents one hamster. Data are from one experiment, n = 5 per condition. *P*-values from two-tailed Mann-Whitney tests (A–C) and unpaired two-tailed *t*-tests (D).

To define the RBD residues targeted by 2C08, we used VSV-SARS-CoV-2-S chimeric viruses (S from D614G strain) to select for variants that escape 2C08 neutralization as previously described (*31*, *33*). We performed plaque assays on Vero cells with 2C08 in the overlay, purified the neutralization-resistant plaques, and sequenced the S genes (**Fig. 3A, Fig. S2A**). Sequence analysis identified the S escape mutations G476D, G476S, G485D, F486P, F486V and N487D, all of which are within the RBD and map to residues involved in ACE2 binding (**Fig. 3B**). To determine whether any of the 2C08 escape mutants we isolated are represented among SARS-CoV-2 variants circulating in humans, we screened all publicly available genome sequences of SARS-CoV-2 (*34*, *35*). Using 829,162 genomes from Global Initiative on Sharing Avian Influenza Data (GISAID), we calculated each substitution frequency in the identified residues site. Of the six escape variants we identified, four were detected among circulating isolates of SARS-CoV-2. The frequency of these substitutions among clinical isolates detected so far is exceedingly rare, with the escape variants representing less than 0.008% of sequenced viruses. In comparison, the D614G substitution is present in 49% of sequenced isolates (**Fig. S2B**).

**Figure 3.**
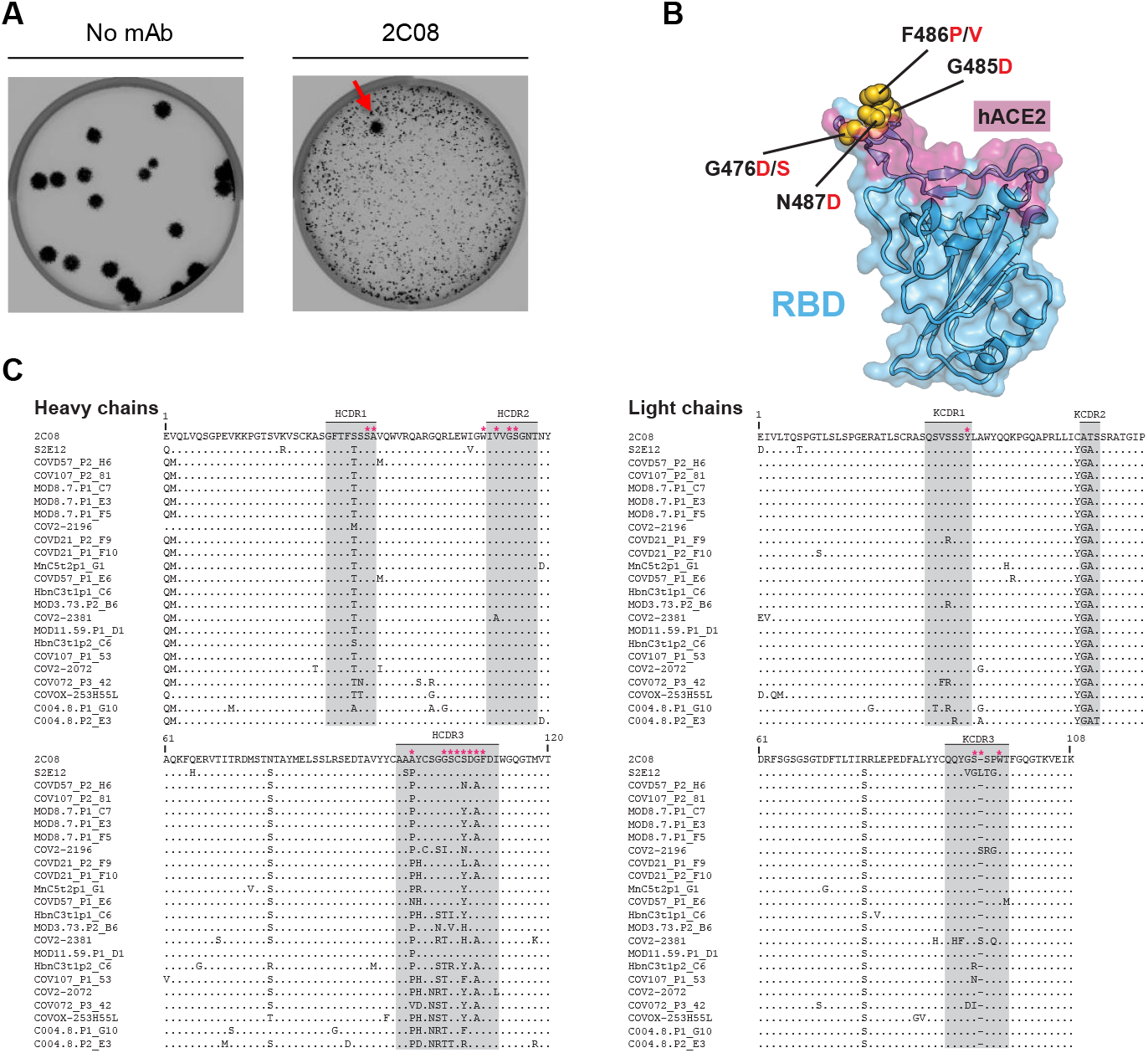
mAb 2C08 recognizes a public epitope in SARS-CoV-2 RBD. (**A**) Plaque assay on Vero cells with no antibody (*left*) or 2C08 (*right*) in the overlay to isolate escape mutants (red arrow). Data are representative of three experiments. (**B**) Structure of RBD (from PDB 6M0J) with hACE2 footprint highlighted in magenta and amino acids whose substitution confers resistance to 2C08 in plaque assays highlighted in yellow. (**C**) Sequence alignment of 2C08 with RBD-binding mAbs from SARS-CoV-2 infected patients and vaccinees that utilize the same immunoglobulin heavy and light chain variable region genes (see also Table S1). Stars indicate contact residues.

We noted that 2C08 targeted residues are similar to those recognized by a previously described human mAb, S2E12, which was isolated from an infected patient (*25*). S2E12 shares a high sequence identity with 2C08 (95% amino acid identity) and is encoded by the same immunoglobulin heavy and light chain variable region genes (**Fig. 3C, Table S1**). Similar to 2C08, S2E12 exhibits potent neutralizing activity *in vitro* and protective capacity *in vivo*. The cryo-EM structure of S2E12 in complex with S shows that the mAb recognizes an RBD epitope that partially overlaps with the ACE2 receptor footprint known as the receptor binding motif (*25*) (**Fig. S3**). S2E12 heavy chain amino acid residues that engage the RBD are identical to those in 2C08, suggesting that 2C08 likely engages the RBD in a manner similar to that of the structurally characterized S2E12. Furthermore, we identified two additional human mAbs, 253H55L and COV2-2196, that share genetic and functional features with 2C08 and have nearly identical antibody-RBD interactions as those of S2E12 (*22*, *36*) (**Fig. 3C, Fig. S3**). Dong et al. noted that COV2-2196 is likely part of a public B cell clone, citing S2E12 and mAbs generated by two other groups which have similar characteristics (*24*, *25*). This prompted us to conduct an expanded search for 2C08-like clonotypes and mAbs. We identified 20 additional mAbs that share the same genetic attributes of 2C08, S2E12, 253H55L and COV2-2196 isolated by different groups from SARS-CoV-2 patients or vaccine recipients (**Fig. 3C and Table S1**). The primary contact residues described for S2E12 were largely conserved for all mAbs (**Fig. 3C**).

Cloning and expression of recombinant human mAbs from single cell sorted B cells is now an established method for generating potential therapeutics against a variety of human pathogens. The source cells are predominantly plasmablasts or memory B cells that are isolated from blood after infection or vaccination (*37*). Here, we describe 2C08, a SARS-CoV-2 vaccine-induced mAb cloned from a GC B cell clone isolated from a draining axillary lymph node sampled from a healthy adult after receiving their second dose of mRNA-based vaccine (*29*). 2C08 is a potently neutralizing antibody that targets the receptor binding motif within the RBD of SARS-CoV-2 S protein and blocks infection by circulating SARS-CoV-2 and emerging variants of concern both *in vitro* and *in vivo*.

2C08 is a “public” mAb, meaning that it is encoded by multiple B cell clonotypes isolated from different individuals that share similar genetic features (*36*). Public antibody responses in humans have been observed after many infections, including SARS-CoV-2 infection (*24*, *36*, *38*–*43*). In the case of 2C08-like clonotypes, the mAbs not only share the immunoglobulin heavy and light chain variable region genes, but also have near identical CDRs and are functionally similar. Several have been shown to bind RBD and neutralize D614G as well as variants B.1.17 and B.1.351. 2C08-like mAbs were isolated from multiple SARS-CoV-2 infected patients independently of demographics or severity of infection. Robbiani *et al*. isolated 2C08-like mAbs from three of six infected individuals analyzed (*24*). Tortorici *et al.* and Zost *et al*. detected a 2C08-like antibody in one or both of two infected individuals they examined, respectively, whereas Kreer *et al*. detected a 2C08-like clone in two of seven patients, in one of whom it was expanded (*22*, *25*, *26*). Wang *et al*. isolated 2C08-like mAbs from five of 14 individuals who received a SARS-CoV-2 mRNA-based vaccine (*15*). Nielsen *et al*. identified 2C08-like rearrangements in sequences derived from four of 13 SARS-CoV-2 patients (*44*). It remains to be determined what fraction of the antibody responses induced by SARS-CoV-2 vaccines in humans are comprised of 2C08-type antibodies that are public, potently neutralizing, and so far, minimally impacted by the mutations found in the variants of concern. It is important to note that at least one 2C08-like mAb, COV2-2196, is currently being developed for clinical use (*36*).

Notably, most of SARS-CoV-2 vaccine induced anti-RBD mAbs also recognized RBDs from the recent variants. It is of some concern, however, that four of the five neutralizing anti-RBD mAbs lost their activity against the B.1.351 and B.1.1.248 SARS-CoV-2 variants. This is consistent with the data reported by Wang *et al*. showing that the neutralizing activity of 14 of 17 vaccine induced anti-RBD mAbs was abolished by the introduction of the mutations associated with these variants (*15*). More extensive analyses with a larger number of mAbs that target the RBD and non-RBD sites will be needed to precisely determine the fraction of vaccine-induced neutralizing antibody response that is compromised due to antigenic changes in emerging SARS-CoV-2 variants of concern. We note that somewhat higher levels of lung viral RNA were recovered from the 2C08-treated animals challenged with the B.1.351-like variant compared to those challenged with the D614G strain. This was unexpected given the similar *in vitro* potency of 2C08 against both viruses and its capacity to protect animals from both groups against weight loss equivalently. One possibility is that 2C08 more readily selected for a partial escape mutant against viruses displaying the B.1.351 variant spike than the WA1/2020 D614G spike.

Given the germinal center B cell origin of 2C08, the binding of 2C08-related clones could be further refined through somatic hypermutation, and their descendants could become part of the high affinity memory B cell and long-lived plasma cell compartments that confer durable protective immunity. Together, these data suggest that first-generation SARS-CoV-2 mRNA-based vaccines can induce public antibodies with robust neutralizing and potentially durable protective activity against ancestral circulating and key emerging SARS-CoV-2 variants.

## Acknowledgements

The Vero-TMPRSS2 cells were kindly provided by Siyuan Ding (Washington University School of Medicine, St. Louis, MO).

## Funding

The contents of this publication are solely the responsibility of the authors and do not necessarily represent the official views of NIAID or NIH. The Ellebedy laboratory was supported by NIAID grant U01AI141990, U01AI150747, NIAID Centers of Excellence for Influenza Research and Surveillance contract HHSN272201400006C. The Ellebedy and Krammer laboratories were supported by NIAID Centers of Excellence for Influenza Research and Surveillance contract HHSN272201400008C, and NIAID Collaborative Influenza Vaccine Innovation Centers contract 75N93019C00051. Work in the Diamond laboratory was partially supported by was supported by NIH contract 75N93019C00062 and grant R01 AI157155. The Shi laboratory was supported by NIH grants AI134907 and UL1TR001439, and awards from the Sealy & Smith Foundation, Kleberg Foundation, the John S. Dunn Foundation, the Amon G. Carter Foundation, the Gilson Longenbaugh Foundation, and the Summerfield Robert Foundation. JST was supported by NIAID 5T32CA009547. JBC was supported by a Helen Hay Whitney postdoctoral fellowship.

## Author contributions

A.J.S. and J.S.T. isolated the antibodies. A.J.S., Z.L., I.D.A., R.E.C., A.J., T.L.B., T.L.D., W.B.A., and J.B.C functionally characterized the mAbs *in vitro* and *in vivo*. T.L., M.T. and F.A. expressed and purified the mAbs and recombinant viral proteins. A.J.S. and J.S.T. compiled and analyzed data. J.A.O and R.M.P. supervised the vaccination study. P.S. and F.K. provided critical reagents. G.B. supervised the structural analysis. S.P.J.W. supervised the mapping analysis. M.S.D. and A.C.M.B. supervised the *in vitro* and *in vivo* analyses, respectively. A.H.E. conceptualized the study and wrote the manuscript with input from all co-authors.

## Competing interests

The Ellebedy laboratory received funding under sponsored research agreements that are unrelated to the data presented in the current study from Emergent BioSolutions and from AbbVie. A.H.E. is a consultant for Mubadala Investment Company. M.S.D. is a consultant for Inbios, Vir Biotechnology, NGM Biopharmaceuticals, Carnival Corporation and on the Scientific Advisory Board of Moderna and Immunome. The Diamond laboratory has received unrelated sponsored research agreements from Moderna, Vir Biotechnology, and Emergent BioSolutions. A patent application related to this work has been filed by Washington University School of Medicine. The Icahn School of Medicine at Mount Sinai has filed patent applications relating to SARS-CoV-2 serological assays and NDV-based SARS-CoV-2 vaccines which list Florian Krammer as co-inventor. Mount Sinai has spun out a company, Kantaro, to market serological tests for SARS-CoV-2. Florian Krammer has consulted for Merck and Pfizer (before 2020), and is currently consulting for Pfizer, Seqirus and Avimex. The Krammer laboratory is also collaborating with Pfizer on animal models of SARS-CoV-2. The Shi laboratory has received sponsored research agreements from Pfizer, Gilead, Merck and IGM Sciences Inc. The Whelan laboratory has received unrelated funding support in sponsored research agreements with Vir Biotechnology, AbbVie and sAB therapeutics. All other authors declare no conflict of interest.

## Data and materials availability

All data are available in the main text or the supplementary materials. Materials used in the analysis are available from the participating laboratories under standard academic material transfer agreements.

## Materials and Methods

### Cell lines

Expi293F cells were cultured in Expi293 Expression Medium (Gibco). Vero-TMPRSS2 cells (a gift from Siyuan Ding, Washington University School of Medicine) were cultured at 37°C in Dulbecco’s Modified Eagle medium (DMEM) supplemented with 10% fetal bovine serum (FBS), 10 mM HEPES pH 7.3, 1 mM sodium pyruvate, 1× non-essential amino acids, and 100 U/ml of penicillin–streptomycin.

### Viruses

The 2019n-CoV/USA_WA1/2020 isolate of SARS-CoV-2 was obtained from the US Centers for Disease Control. The UK B.1.1.7 isolate was obtained from an infected individual (*16*). The point mutation D614G in the spike gene was introduced into an infectious complementary DNA clone of the 2019n-CoV/USA_WA1/2020 strain as described previously (*45*). The generation of a SARS-CoV-2 virus with the South African variant spike gene (B.1.351) in the background of 2019n-CoV/USA_WA1/2020 was described previously (*16*). All viruses were passaged once in Vero-TMPRSS2 cells and subjected to deep sequencing after RNA extraction to confirm the introduction and stability of substitutions (*16*). All virus preparation and experiments were performed in an approved Biosafety level 3 (BSL-3) facility.

### Monoclonal antibody (mAb) generation

Antibodies were cloned as described previously (*46*). Briefly, VH, Vκ, and Vλ genes were amplified by reverse transcription-PCR and nested PCR reactions from singly sorted GC B cells using primer combinations specific for IgG, IgM/A, Igκ, and Igλ from previously described primer sets (*47*) and then sequenced. To generate recombinant mAbs, restriction sites were incorporated via PCR with primers to the corresponding heavy and light chain V and J genes. The amplified VH, Vκ, and Vλ genes were cloned into IgG1, Igκ, and Igλ expression vectors, respectively, as described previously (*47*–*49*). Heavy and light chain plasmids were co-transfected into Expi293F cells (Gibco) for expression, and mAbs were purified with protein A agarose (GoldBio).

### Antigens

Recombinant receptor binding domain of S (RBD), was expressed as previously described (*50*, *51*). Briefly, RBD, along with the signal peptide (amino acids 1-14) plus a hexahistidine tag were cloned into mammalian expression vector pCAGGS. RBD mutants were generated in the pCAGGS RBD construct by changing single residues using mutagenesis primers. Recombinant proteins were produced in Expi293F cells (ThermoFisher) by transfection with purified DNA using the ExpiFectamine 293 Transfection Kit (ThermoFisher). Supernatants from transfected cells were harvested 4 days post-transfection, and recombinant proteins were purified using Ni-NTA agarose (ThermoFisher), then buffer exchanged into phosphate buffered saline (PBS) and concentrated using Amicon Ultracel centrifugal filters (EMD Millipore).

### Enzyme-linked immunosorbant assay

Assays were performed in 96-well plates (MaxiSorp; Thermo). Each well was coated with 100 μL of wild-type or variant RBD or bovine serum albumin (1μg/mL) in PBS, and plates were incubated at 4 °C overnight. Plates were then blocked with 0.05% Tween20 and 10% FBS in PBS. mAbs were serially diluted in blocking buffer and added to the plates. Plates were incubated for 90 min at room temperature and then washed 3 times with 0.05% Tween-20 in PBS. Goat anti-human IgG-HRP (Jackson ImmunoResearch 109-035-088, 1:2,500) was diluted in blocking buffer before adding to wells and incubating for 60 min at room temperature. Plates were washed 3 times with 0.05% Tween20 in PBS, and then washed 3 times with PBS. o-Phenylenediamine dihydrochloride substrate dissolved in phosphate-citrate buffer (Sigma-Aldrich) with H_2_O_2_ catalyst was incubated in the wells until reactions were stopped by the addition of 1 M HCl. Optical density measurements were taken at 490 nm. Area under the curve was calculated using Graphpad Prism v8.

### Focus reduction neutralization test

Serial dilutions of each mAb diluted in DMEM with 2% FBS were incubated with 10_2_ focus-forming units (FFU) of different strains or variants of SARS-CoV-2 for 1 h at 37°C. Antibody-virus complexes were added to Vero-TMPRSS2 cell monolayers in 96-well plates and incubated at 37°C for 1 h. Subsequently, cells were overlaid with 1% (w/v) methylcellulose in MEM supplemented with 2% FBS. Plates were harvested 24 h later by removing overlays and fixed with 4% PFA in PBS for 20 min at room temperature. Plates were washed and sequentially incubated with an oligoclonal pool of SARS2-2, SARS2-11, SARS2-16, SARS2-31, SARS2-38, SARS2-57, and SARS2-71 anti-S antibodies (*33*) and HRP-conjugated goat anti-mouse IgG (Sigma 12-349) in PBS supplemented with 0.1% saponin and 0.1% bovine serum albumin. SARS-CoV-2-infected cell foci were visualized using TrueBlue peroxidase substrate (KPL) and quantitated on an ImmunoSpot microanalyzer (Cellular Technologies).

### SARS-CoV-2 hamster studies

All procedures involving animals were performed in accordance with guidelines of the Institutional Animal Care and Use Committee of Washington University in Saint Louis. Four- to six-week old male Syrian hamsters were obtained from Charles River Laboratories and housed in an enhanced ABSL3 facility at Washington University in St Louis. Animals were randomized from different litters into experimental groups and were acclimatized at the BSL3 facilities for 4-6 days prior to experiments. Animals received intra-peritoneal (IP) injection of isotype control or anti-SARS-CoV-2 mAbs 24 h prior to SARS-CoV-2 challenge. Hamsters were anesthetized with ketamine (150 mg/kg) and xylazine (10 mg/kg) via IP injection and were intranasally inoculated 5 × 10^5^ PFU of 2019n-CoV/USA_WA1/2020-D614G or Wash SA-B.1.351 SARS-CoV-2 in 100 μL PBS. Animal weights were measured every day for the duration of experiments. Animals were euthanized 4 dpi and the lungs were collected for virological analyses. Left lung lobes were homogenized in 1 mL of PBS or DMEM, clarified by centrifugation, and used for virus titer and cytokine assays.

### Virus titration assays from hamster lung homogenates

Plaque assays were performed on Vero-Creanga cells in 24-well plates. Lung tissue homogenates were serially diluted 10-fold, starting at 1:10, in cell infection medium (DMEM + 2% FBS + L-glutamine + penicillin + streptomycin). Two hundred and fifty microliters of the diluted virus were added to a single well per dilution per sample. After 1 h at 37°C, the inoculum was aspirated, the cells were washed with PBS, and a 1% methylcellulose overlay in MEM supplemented with 2% FBS was added. Seventy-two hours after virus inoculation, the cells were fixed with 4% formalin, and the monolayer was stained with crystal violet (0.5% w/v in 25% methanol in water) for 1 h at 20°C. The number of plaques were counted and used to calculate the plaque forming units/mL (PFU/mL).

To quantify viral load in lung tissue homogenates, RNA was extracted from 140 μL samples using QIAamp viral RNA mini kit (Qiagen) and eluted with 50 μL of water. Four μL RNA was used for real-time qRT-PCR to detect and quantify N gene of SARS-CoV-2 using TaqMan™ Fast Virus 1-Step Master Mix as described (*53*) or using the following primers and probes: Forward: GACCCCAAAATCAGCGAAAT; Reverse: TCTGGTTACTGCCAGTTGAATCTG; Probe: ACCCCGCATTACGTTTGGTGGACC; 5’Dye/3’Quencher: 6-FAM/ZEN/IBFQ. Viral RNA was expressed as (N) gene copy numbers per mg for lung tissue homogenates, based on a standard included in the assay, which was created via *in vitro* transcription of a synthetic DNA molecule containing the target region of the N gene.

### Cytokine assay from hamster lung homogenates

RNA extracted from hamster lung homogenates was reverse transcribed using random hexamers and Superscript III (Thermo Scientific) with the addition of RNase inhibitor according to the manufacturer’s protocol. Gene expression of inflammatory cytokines was determined using PrimeTime Gene Expression Master Mix (Integrated DNA Technologies) with primer/probe sets specific for RPL18 (Accession number: XM_005084699.3, Forward: GTTTATGAGTCGCACTAACCG, Reverse: TGTTCTCTCGGCCAGGAA, Probe: TCTGTCCCTGTCCCGGATGATC) (*54*); CCL3 (Accession number: NM_001281338.1, Forward: CCTCCTGCTGCTTCTTCTATG, Reverse: TGCCGGTTTCTCTTGGTTAG, Probe: TCCCGCAAATTCATCGCCGACTAT); CCL5 (Accession number: XM_005076936.3, Forward: TGCTTTGACTACCTCTCCTTTAC, Reverse: GGTTCCTTCGGGTGACAAA, Probe: TGCCTCGTGTTCACATCAAGGAGT); IFIT3 (Accession number: XM_021224964.1, Forward: CTGATACCAACTGAGACTCCTG, Reverse: CTTCTGTCCTTCCTCGGATTAG, Probe: ACCGTACAGTCCACACCCAACTTT); IL-6 (Accession number: XM_005087110.2, Forward: CCAGATCTACCTGGAGTTTGTG, Reverse: CTGGACCCTTTACCTCTTGTTT, Probe: AAGCCAGAGTCATTCAGAGCACCA); IP10 (Accession number: NM_001281344.1, Forward: AGAGCCTCTTAACCAGAGAGAA, Reverse: TAGCCATAGGCCGACGTATAA, Probe: AAAGCCCGTCTCTCCATCACTTCT); IRF7 (Accession number: XM_005063345.3, Forward: AGCACGGGACGCTTTATC, Reverse: GACGGTCACTTCTTCCCTATTC, Probe: AGTTTGGATGTACTGAAGGCCCGG); RIG-I (Accession number: NM 001310553.1, Forward: GTGCAACCTGGTCATTCTTTATG, Reverse: GTCAGGAGGAAGCACTTACTATC, Probe: AAACCAGAGGCAGAGGAAGAGCAA).

Cytokine expression levels were compared between 2019n-CoV/USA_WA1/2020-D614G or Wash SA-B.1.351 SARS-CoV-2 challenged hamsters and naïve controls using the −ΔΔCt method.

### Selection of 2C08 mAb escape mutants in SARS-CoV-2 S

We used VSV-SARS-CoV-2-S chimera with S D614G to select for SARS-CoV-2 S variants that escape mAb neutralization described as previously (*31*, *33*). Antibody neutralization resistant mutants were recovered by plaque isolation. Briefly, plaque assays were performed to isolate the VSV-SARS-CoV-2 escape mutant on Vero cells with mAb 2C08 in the overlay. The concentration of 2C08 in the overlay was determined by neutralization assays at a multiplicity of infection (MOI) of 100. Escape clones were plaque-purified on Vero cells in the presence of 2C08, and plaques in agarose plugs were amplified on MA104 cells with the 2C08 present in the medium. Viral supernatants were harvested upon extensive cytopathic effect and clarified of cell debris by centrifugation at 1,000 × g for 5 min. Aliquots were maintained at −80°C. Viral RNA was extracted from VSV-SARS-CoV-2 mutant viruses using RNeasy Mini kit (Qiagen), and S was amplified using OneStep RT-PCR Kit (Qiagen). The mutations were identified by Sanger sequencing (GENEWIZ).

**Figure S1.**
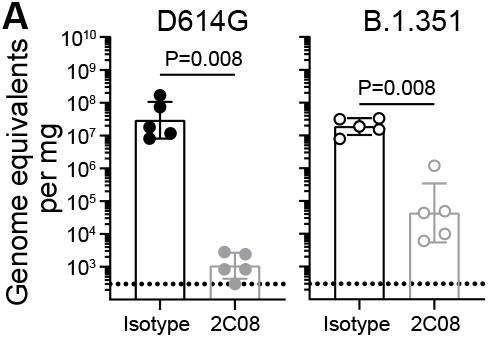
mAb 2C08 protects hamsters from SARS-CoV-2 challenge. (**A**) Lung viral RNA titer using 5’ UTR probe of hamsters that received isotype (black) or 2C08 (grey) one day prior to intranasal challenge with 10^5^ TCID_50_ D614G (*left*) or B.1.351 (*right*) SARS-CoV-2 variants. Bars indicate geometric mean ± geometric SD, and each symbol represents one hamster. Data are from one experiment, n = 5 per condition. *P*-values from two-tailed Mann-Whitney tests.

**Figure S2.**
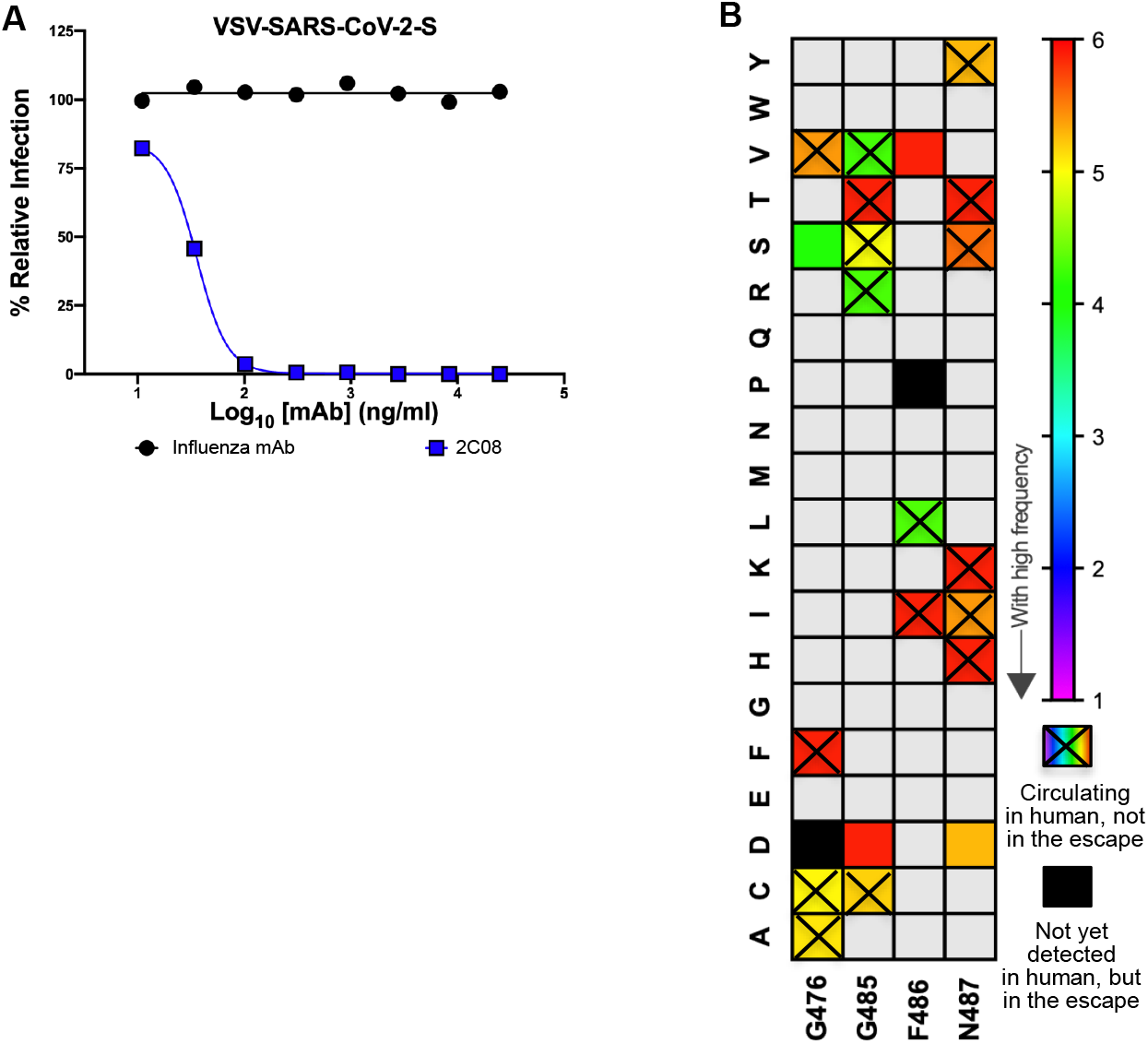
Escape mutant mapping of mAb 2C08. (**A**) 2C08 and a control anti-influenza virus mAb were tested for neutralizing activity against VSV-SARS-CoV-2. The concentration of 2C08 added in the overlay completely inhibited viral infection. Data are representative of two independent experiments. (**B**) 2C08 escape profile in currently circulating SARS-CoV-2 viruses isolated from humans. For each site of escape, we counted the sequences in GISAID with an amino acid change (829,521 total sequences at the time of the analysis). Variant circulating frequency is represented as a rainbow color map from red (less circulating with low frequency) to violet (most circulating with high frequency). A black cell indicates the variant has not yet been isolated from a patient. A rainbow cell with cross indicates the variant has been isolated from a patient, but not appear in those 2C08 mAb escape mutants.

**Figure S3.**
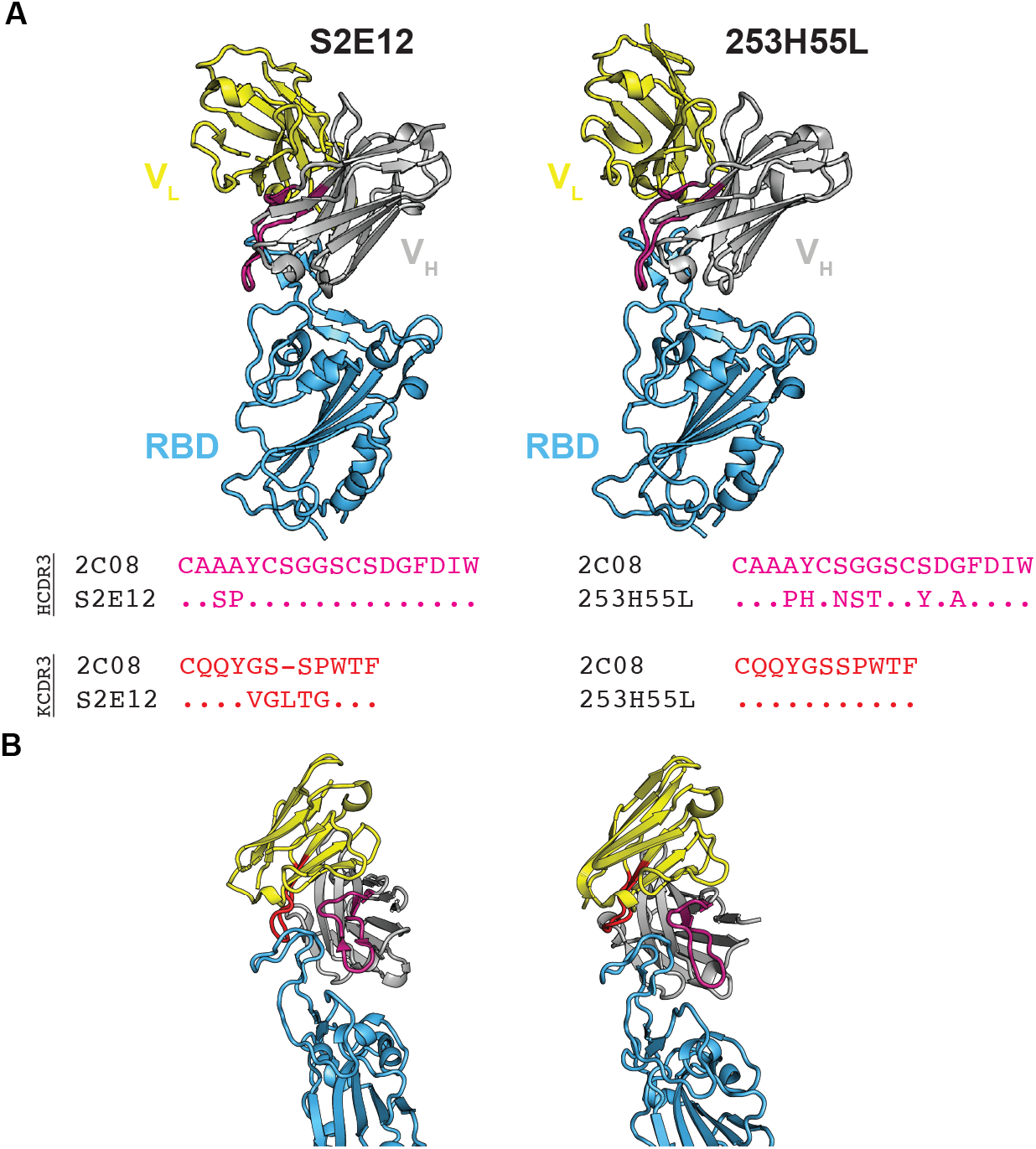
mAb 2C08 recognizes a public epitope in SARS-CoV-2 RBD. (**A and B**) Structures of mAbs S2E12 (PDB 7K45) and 253H55L (PDB 7ND9) complexed with RBD and their heavy (pink) and light (red) chain CDR3 sequence alignments with 2C08.

**Table S1.**
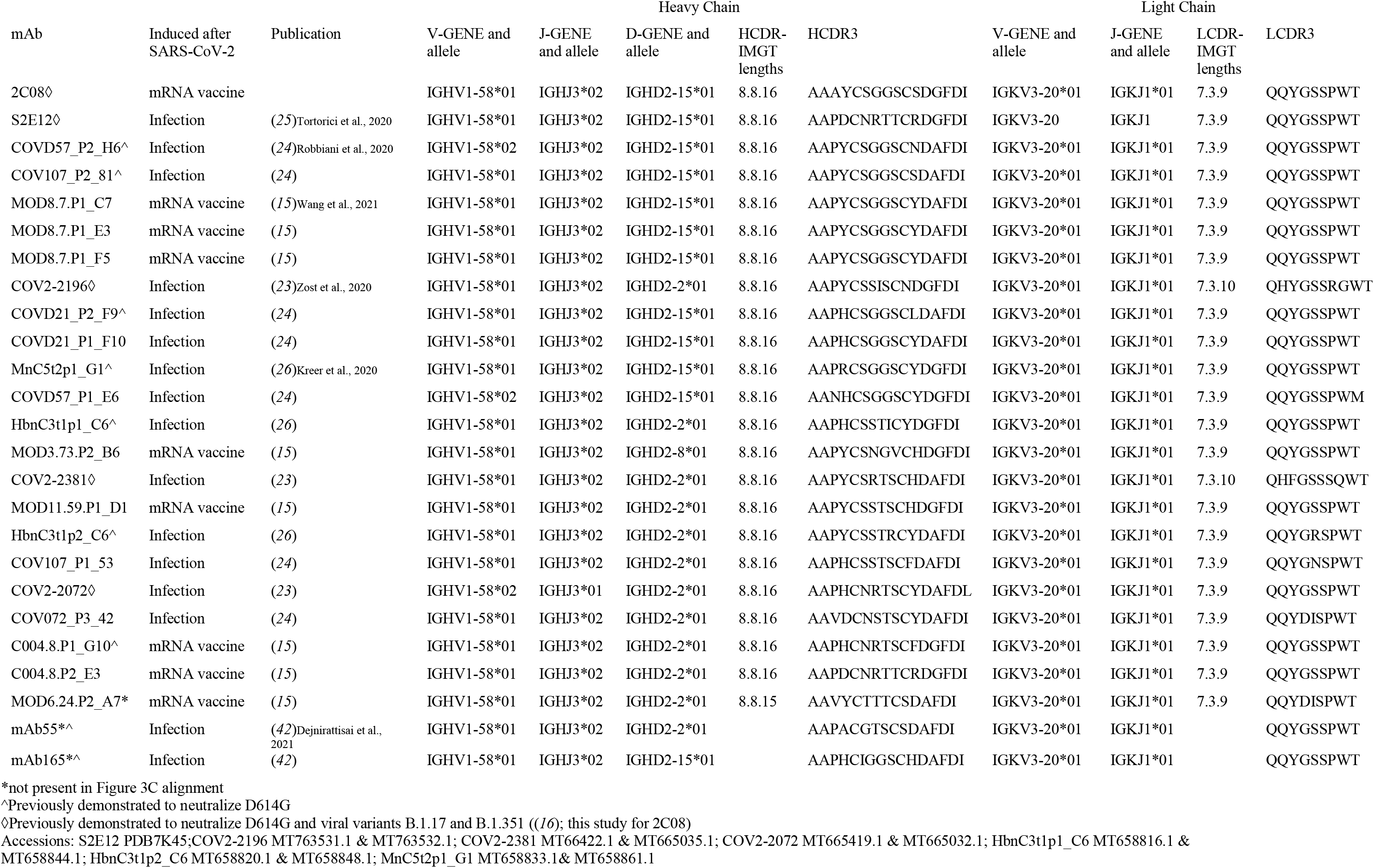

